# Differences in neuronal numbers, morphology and developmental apoptosis in mice nigra provide experimental evidence of ontogenic origin of vulnerability to Parkinson’s disease

**DOI:** 10.1101/2020.12.10.419259

**Authors:** D J Vidyadhara, Haorei Yarreiphang, Trichur R Raju, Phalguni Anand Alladi

## Abstract

Parkinson disease (PD) prevalence varies by ethnicity. In an earlier study we replicated the reduced vulnerability to PD in an admixed population, using 1-methyl-4-phenyl-1,2,3,6-tetrahydropyridine (MPTP)-susceptible C57BL/6J, MPTP-resistant CD-1 and their F1 crossbreds. In the present study we investigated if the differences have a developmental origin. Substantia nigra was evaluated at postnatal days 2 (P2), P6, P10, P14, P18, and P22. C57BL/6J mice had smaller nigra and fewer dopaminergic neurons than the CD-1 and crossbreds at P2, which persisted through development. A significant increase in numbers and nigral volume was observed across strains till P14. A drastic decline thereafter was specific to C57BL/6J. CD-1 and crossbreds retained their numbers from P14 to stabilize with supernumerary neurons at adulthood. The neuronal size increased gradually to attain adult morphology at P10 in the resistant strains, vis-à-vis at P22 in C57BL/6J. Accordingly, in comparison to C57BL/6J, the nigra of CD-1 and reciprocal crossbreds possessed cyto-morphological features of resilience, since birth. The considerably lesser dopaminergic neuronal loss in the CD-1 and crossbreds seen at P2, P14 and thereafter was complemented by attenuated developmental cell death. The differences in programmed cell death were confirmed by reduced TUNEL labelling, AIF and caspase-3 expression. GDNF expression aligned with the cell death pattern at P2 and P14 in both nigra and striatum. Earlier maturity of nigra and its neurons appear to be better features that reflect as MPTP-resistance at adulthood. Thus variable MPTP-vulnerability in mice and also differential susceptibility to PD in humans may arise early during nigral development.

## Introduction

Degeneration of A9 subset of dopaminergic (DA) neurons of the substantia nigra pars compacta (SNpc) is a key neuropathological hallmark of Parkinson’s disease (PD). A loss of 50-60% DA neurons is a pre-requisite to manifest the motor abnormalities (Fearnley & Lees 1991; Ma *et al.* 1996; Kordower *et al.* 2013; Vidyadhara *et al.* 2019a) which points at the requirement of a critical numbers of DA neurons for normal function or setting off the pathology. In this context, Fahn (1989) hypothesised that individuals born with lesser number of nigral neurons are more likely to develop PD, which raises a possibility that quantitative differences between nigral neuronal numbers could be the basis for ethno-racial bias in PD prevalence. USA-based Caucasians have high prevalence of PD (329.3/ 100,000) (Strickland & Bertoni 2004) along with the European-Caucasians (108-257/100,000) (von Campenhausen *et al.* 2005). A recent study on North American population revealed an alarming rise in PD prevalence to 572/100,000 (Marras et al., 2018). A Chicago-based study documented age-related loss of nigral DA phenotype (Chu *et al.* 2002), while other Europe-based studies showed frank age-related neuronal loss (Ma *et al.* 1996; Cabello *et al.* 2002). In contrast, Asian-Indians with lower PD prevalence (52.7/100,000) (Das *et al.* 2010) had relatively higher number of SNpc DA neurons with no age-related neuronal loss (Alladi *et al.* 2009). This along with subtle glial; dendritic and synaptic changes noted in this population (Naskar *et al.* 2019; Jyothi *et al.* 2015) might explain the lower prevalence of PD in them. Thus DA neuronal numbers and their response to aging may play a vital role in shaping the individual’s vulnerability to develop PD. The prevalence of PD in admixed populations is of potential interest due to the complex genetic patterns resulting from admixing and the likelihood of alterations in the disease patterns. The Anglo-Indians who are the first generation admixed population born to an Asian-Indian female and a Caucasian male are, surprisingly, at a much lesser risk for PD (Ragothaman *et al.* 2003). In the absence of autopsied tissues of this population, we designed an animal model of admixing by studying the F1 generation of 1-methyl-4-phenyl-1,2,3,6-tetrahydropyridine (MPTP)-resistant CD-1 mated with MPTP-susceptible C57BL/6J strains of mice. Our model effectively recapitulated the “much lesser risk for PD” seen in the Anglo-Indians citing the differences in cyto-morphological patterns, explaining the differential prevalence of PD to a considerable extent (Vidyadhara *et al.* 2017). In brief, the C57BL/6J mice, which possessed fewer DA neurons, were highly vulnerable to MPTP, compared to CD-1, which had more of these neurons. Other DA phenotypes such as cellular size along with tyrosine hydroxylase (TH), Nurr1, PitX3, calbindin D28k, glial derived neurotrophic factor, caspase-3, mitochondrial fission/fusion proteins and interneuron related proteins also appeared to influence the MPTP-induced susceptibility/resistance (Vidyadhara *et al.* 2016; Vidyadhara *et al.* 2019b; Seshadri & Alladi 2019; Bhaduri *et al.* 2018; Vidyadhara *et al.* 2017). The F1 crossbreds of these strains were much better protected against MPTP, with the particularly striking finding of supernumerary nigral DA neurons and absence of neuronal loss in response to MPTP-toxicity. Preserved motor behavior and compensated/unaffected striatal local field potentials in PD reminiscent conditions in CD-1 and the crossbreds complement the cytomolecular observations (Vidyadhara *et al.* 2019b). These features indicate that admixing imparts resilience.

Cytomorphological and molecular features of maturation of the nigral DA neurons are seeded during ontogenesis (Sailaja & Gopinath 1996; Sailaja & Gopinath 1994). The total number of DA neurons at adulthood may thus be finalised during the postnatal period. Aspects like establishment of DA phenotype, synaptogenesis, trophic factor supply, levels of pro-apoptotic and anti-apoptotic proteins and other cytomolecular attributes during this period are critical determinants of their survival (Jackson-Lewis *et al.* 2000). Study of substantia nigra DA neurons in different ethnicities during the formative period might reveal the mechanisms, however, due to the valid ethical constraints that limit the availability of human foetal and neonatal brain specimens, it is difficult to conduct such experiments.

In the fetal primate brains, apoptosis of DA neurons was observed midway through gestation i.e. embryonic day 80 and identified histologically by chromatin clumping in tyrosine hydroxylase-positive cells and confirmed by Terminal deoxynucleotidyl transferase (TdT) dUTP Nick-End Labeling (TUNEL) and caspase-3 staining (Morrow et al., 2007), with a loss of at least 50% of the DA neurons. In developing rats, programmed cell death (PCD) in the midbrain DA system began at the end of gestation to peak in the postnatal period (Oo & Burke, 1997). Caspase-3 and apoptosis inducing factor (AIF) are accepted modulators of apoptotic cascades in the developing nigra. GDNF appears to be the most potent supporter of embryonic nigral DA neurons (Burke, 1998). Disruption of nuclear membrane integrity and release of pro-aggregant nuclear factors, such as histones, that may trigger α-synuclein aggregation which leads to cell death (Jiang et al., 2016, 2017). Developmental PCD is an important determinant of the size and function of a neuronal population at adulthood and its timing and magnitude is critical to understand the intrinsic and extrinsic factors that affect PCD in DA neurons.

In the present study, we evaluated the postnatal development of SNpc in our model of differential susceptibility using four mice strains i.e. C57BL/6J, CD-1 and their reciprocal crossbred strains, F1X1 and F1X2 (Vidyadhara *et al.* 2019b; Vidyadhara *et al.* 2017). We performed stereological quantification to estimate the number of nigral neurons followed by the evaluation of morphological alterations and compared them with the features seen in adults. These estimations were conducted over the period of postnatal development i.e. P2 (postnatal day 2) to P22 on every 4th day, in order to earmark the critical developmental time points of 1) maximal cell loss 2) morphological maturation and 3) alterations in expression profile of tyrosine hydroxylase (TH). Apoptosis was evaluated by TUNEL assay. AIF and caspase-3 were evaluated by immunofluorescence and confocal microscopy-based evaluation at periods of peak apoptosis i.e. P2, P14 and P22 in SNpc. GDNF expression was evaluated at P2 and P14 in the nigro-striatum. Our study provides experimental evidence that developmental apoptosis sculpts the nigra in terms of neuronal numbers and facilitates the identification of strain-specific critical window of maximum susceptibility of DA neurons.

## Materials and Methods

### Mice strains

The pregnant mice were housed in polypropylene cages under standard laboratory conditions with ad-libitum access to food and water. Age and gender matched neonates (n=6/group/experiment) born following mating of C57BL/6J female and CD-1 males were named F1X1, while the reciprocal ones were named F1X2 (Vidyadhara *et al.* 2017). The experiments were performed in the light period (08:00-18:00h) in accordance with the guidelines of the Committee for the Purpose of Control and Supervision of Experiments on Animals (CPCSEA), New Delhi, India, that are based on NIH, USA guidelines.

### Tissue processing for immunohistochemistry and TUNEL

We anaesthetized neonates aged postnatal day 2 (P2), P6, P10, P14, P18 and P22 of each strain (n=6/strain/time-point) by halothane inhalation and perfused intracardially with saline followed by 4% buffered paraformaldehyde (0.1M phosphate buffer; pH 7.4). The brains were quickly harvested and post-fixed for 24-48h at 4°C. Cryoprotected midbrains (10%, 20%, and 30% sucrose in 0.1M PB) were cryosectioned (Leica Microsystems, Germany) at 40μm. Serial sections were collected on gelatinized slides. Every third midbrain section from P2, P6, and P10 specimens and every sixth midbrain section from P14, P18, and P22 were immunostained, a slight modification of earlier reports (Jackson-Lewis *et al.* 2000).

### Immuno-peroxidase staining of tyrosine hydroxylase (TH)

Briefly, following the quenching of endogenous peroxidase by 0.1% H_2_O_2_ in 70% methanol, the non-specific staining was blocked with 3% buffered bovine serum albumin (BSA). The sections were sequentially incubated with the rabbit polyclonal anti-TH antibody (1:500, SC-25269, Santacruz Biotechnology Inc, USA;72 hr), anti-rabbit secondary antibody (1:200 dilution; Vector Laboratories, Burlingame, USA; 18hr) and avidin–biotin complex (1:100, Elite ABC kits; Vector Laboratories; USA; 4hr). The chromogen was a 0.05% solution of 3’-3’-diaminobenzidine and 0.1M acetate imidazole buffer (pH 7.4) with 0.1% H_2_O_2_. The dilution and washing, were done with 0.01 M PBS containing 0.1% Triton X-100 (0.01M PBST, pH 7.4). In the negative controls, the primary antibody was replaced with the dilution buffer (Suresh *et al.* 2017; Sn *et al.* 2019; Vidyadhara *et al.* 2017).

### Stereology based analysis

Stereological quantification of TH-immunoreactive (TH-ir) DA neurons was performed using optical fractionator probe (Baquet *et al.* 2009; Suresh *et al.* 2018; Vidyadhara *et al.* 2017). On every selected midbrain section, the SN was delineated using a 4X objective of Olympus BX61 Microscope (Olympus Microscopes, Japan) equipped with StereoInvestigator (Version 8.1, Micro-brightfield Inc., Colchester, USA). We counted the TH-ir cells using high power objective (100X), with a regular grid interval of 22500μm^2^ (x=150μm, y=150μm) and counting frame of size 3600μm2 (x=60μm, y=60μm). A guard zone of 4μm was implied on both sides resulting in an optical dissector of 17μm. Quantification was performed in both hemispheres and pooled to derive total numbers.

The nigral volume was measured by principles of contour planimetry by delineating the nigra with 10X objective of Olympus BX61 Microscope (Olympus Microscopes, Japan) during stereological quantification. Evaluation interval, section thickness and grid spacing were identical to that of the parameters for stereological estimates of neuronal numbers. Besides, we qualitatively evaluated the ventral midbrain by light microscopy (4X, Leica DM 750) at different time points, to derive the developmental time-point of attaining adult architecture.

### Densitometry based image analysis and Morphometry

High magnification (40X) images non-overlapping images of TH-ir DA neurons of SNpc were analysed offline for cellular TH expression using Q Win V3 (Leica Systems, Germany). A cumulative mean was derived after sampling approximately 150-200 DA neurons/animal. The values were expressed on an optical density scale of 0-255, where ‘255’ equalled intense staining. Similarly for soma size, around 150 neurons per animal were sampled for P2-P10, and around 200 cells for P14-P22. The cumulative values were expressed in micrometre square (Vidyadhara *et al.* 2017).

### TUNEL and TH co-labelling assay

We used a commercially available TUNEL kit (4823-30-K, Trivigen, USA) as per manufacturer’s instructions, to detect apoptotic nuclei. Briefly, hydrated tissue sections were equilibrated with NeuroPore^TM^ solution (30 min), followed by washing and immersion in TdT labeling buffer. The sections were incubated with labelling reaction mix in a humid chamber (2hr) following which the reaction was terminated with Tdt stop buffer. The sections were then labelled with Strep-HRP solution. The staining was visualized using 0.05% solution of 3’-3’-diaminobenzidine (DAB) in 0.1M acetate imidazole buffer; pH 7.4 and 0.1% H_2_O_2_ with nickel ammonium sulfate enhancement. Thereafter, the sections were stained for TH, by immunoperoxidase protocol, to confirm that the assessment of apoptosis was being performed within the SNpc. (Vidyadhara et al., 2016b). Finally, the sections were dehydrated, cleared, and cover-slipped using DPX. PBST (0.01 M, pH 7.4) was used as both working and washing buffer. For negative controls, the primary antibody was omitted.

### Immunofluorescence staining

A sequential immunostaining protocol was performed to study the expression of AIF (1:200, SC-9416, Santa Cruz Biotechnology Inc, USA) and caspase-3 (1:200, SC-7148, Santa Cruz Biotechnology Inc, USA) as well as GDNF (1:200, SC-328, Santa Cruz Biotechnology Inc, USA) co-labelling with TH (Vidyadhara et al., 2017, Alladi, et al., 2010a). Briefly, the sections were equilibrated (0.1M PBS, pH 7.4) and blocked for 1h in 3% BSA. Thereafter, the sections were incubated with the primary antibody (48 hr at 4°C). This was followed by incubation with appropriate secondary antibodies i.e. Cy3-conjugated (1:200; C2821 or C2181; Sigma Aldrich, USA), FITC-conjugated (1:200; F7367 or F7512, Sigma Aldrich, USA) or Cy5-conjugated secondary antibodies (1:200; AP192SA6; Merck Millipore, USA; 17 hrs at 4°C). Similar immunolabeling steps were followed for the subsequent sequential staining. We used 0.01 M PBST (pH 7.4) as both working and washing buffer. Negative controls sections were incubated with only dilution buffer. Sections were mounted using Vectashield hard set mounting medium (H-400; Vector Laboratories, USA).

### Quantification of immunofluorescence intensity by image analysis

The immunofluorescence-labelled images of different sub-regions of SNpc were captured using laser scanning confocal microscope with the 20X objective at an optical zoom of two (DMIRE-TCS Leica, Germany). The intensity levels were used as measures of protein expression, using the inbuilt software (Alladi et al., 2010b; Vidyadhara, et al., 2016). Around 150-200 neurons/animal were sampled to derive cumulative mean. The excitation frequencies used were 488nm for FITC, 514nm for Cy3 and 633nm for Cy5. Emission band widths of 495–540nm for FITC, 550–620nm for Cy3, and 649-666 nm for Cy5 were maintained to avoid any overlap of emission frequencies. The striatum was photographed with 10X objective at a constant PMT voltage of 537 (Alladi, et al., 2010a & b).

### Statistical Analysis

The data was analysed using two-way ANOVA followed by Tukey’s post hoc test. The values were stated as mean + SD and a p-value lower than 0.05 was considered significant.

## Results

### Maturation of nigral architecture

The SNpc and pars reticulata (SNpr) were indistinguishable across the strains at P2 (Fig. 1 a1-d1). The DA neurons were randomly distributed throughout the ventral midbrain and occasional clusters were noted along the midline between P2 and P6. A gradual distinction between the two tiers was achieved after P6 (Fig. 1, a2, - d2). By P10 the SNpc DA neurons were identifiable (Fig. 1 a3-d3). A clear adult architecture i.e. a compact band of DA neurons rich SNpc was observed at P14 in CD-1 (Fig. 1 b4) and the crossbreds (Fig. c4-d4), whereas this discrimination was relatively delayed in C57BL/6J and appeared at P18 (Fig. 1 a4). Qualitative observations of different nigral subsections showed that, the DA neurons in the lateral nigra appeared at P14 in C57BL/6J compared to appearance at P10 in the other strains (Fig. 1 b3-a4, arrows).

**Fig. 1.**
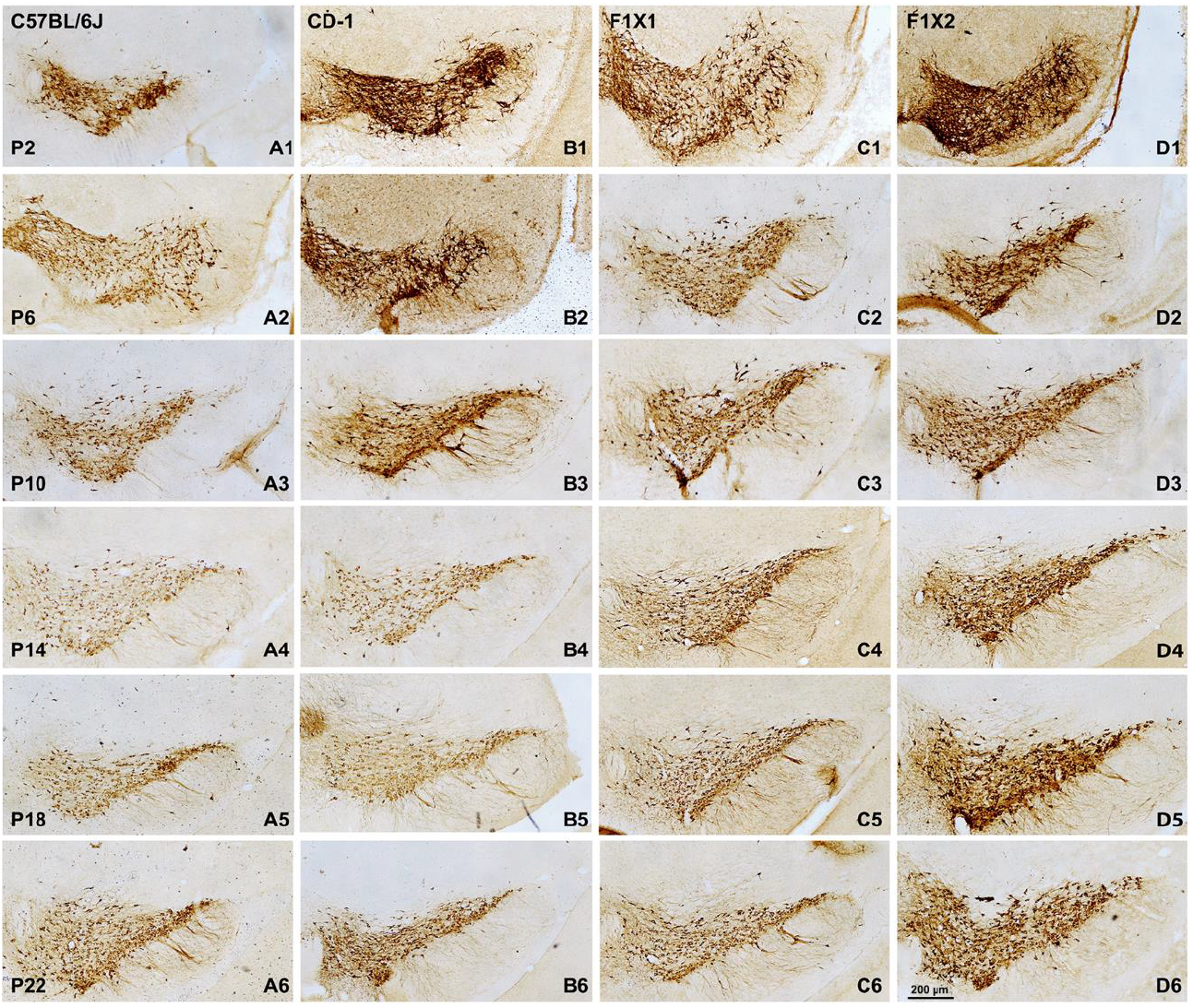
Representative photomicrographs showing that nigral architecture matures earlier in MPTP resistant strains and they have more neurons: The SNpc and pars reticulata (SNpr) are largely indistinguishable at P2 (a1 - d1). Note that, the dopaminergic neurons in the lateral nigra appeared at P6 in the other strains but at P10 in C57BL/6J (a3). By P10 the SNpc appear to be organized (a3-d3). Adult architecture of SNpc is evident at P14 in CD-1 (b4) and the crossbreds (c4-d4), but at P18 in C57BL/6J (a4).

### Differences in numbers of DA neurons during postnatal development

The C57BL/6J mice had significantly fewer TH -IR nigral neurons when compared to CD-1 and the crossbreds at P2 (Fig. 2a, Table 1, P2, ^$$$$^p<0.0001 C57BL/6J v/s CD-1, F1X1 & F1X2). The numbers increased significantly thereafter in all the strains and the maximal increase was noted from P6 to P14 (Fig. 2a). During all the stages studies, the differences between C57BL/6J and rest of the strains were notable.

**Fig. 2.**
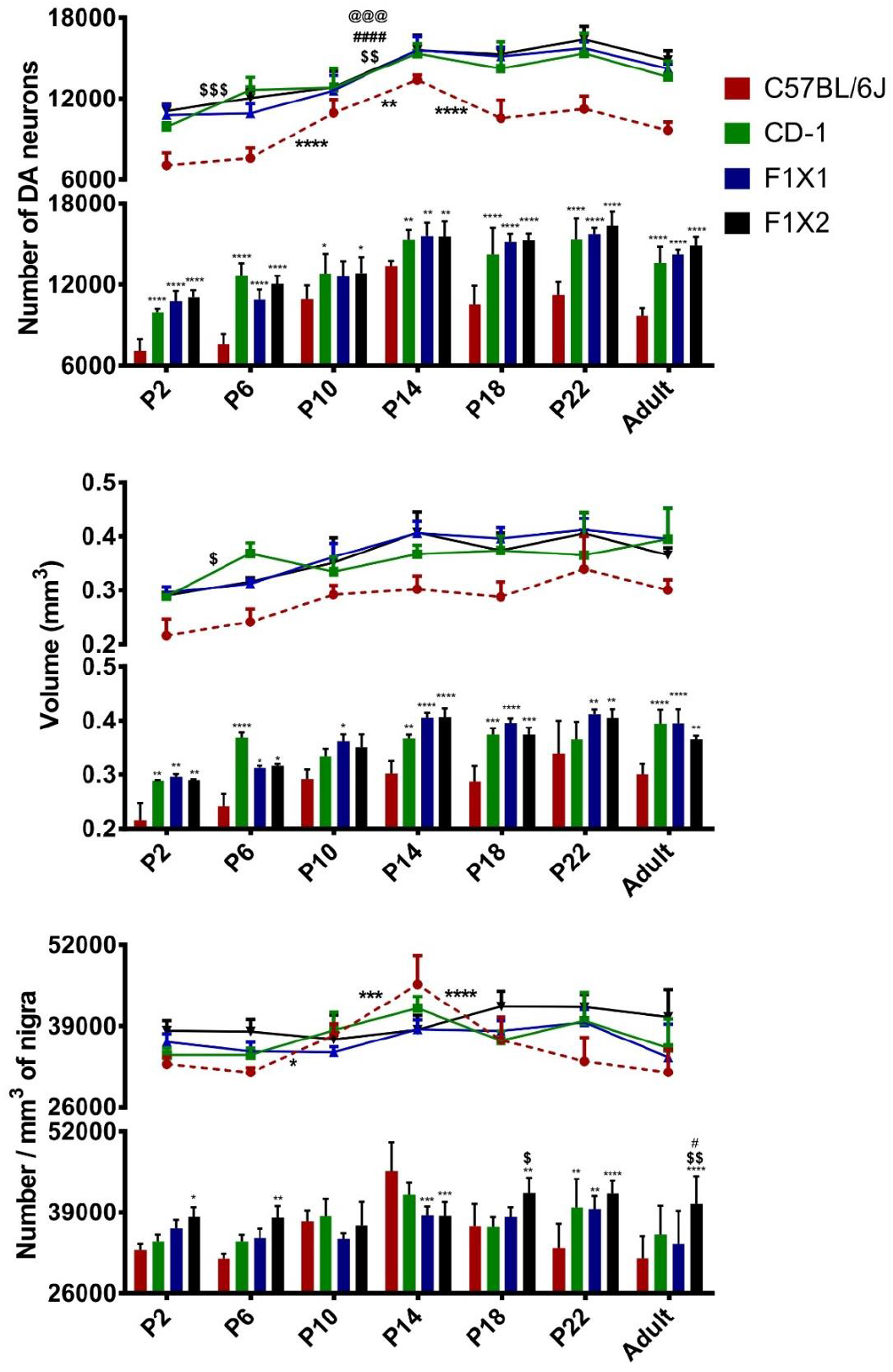
Histograms and line graphs showing alterations in nigral neuronal numbers (a) volume (b) and neuronal density (c). (a) The C57BL/6J mice had significantly fewer nigral neurons when compared to CD-1 and the crossbreds at P2 (P2, ^$$$$^p<0.0001 C57BL/6J v/s CD-1, F1X1 & F1X2). The numbers increased significantly thereafter in all the strains. The period of maximum increase was between P6 to P14. Note the reduction in numbers till P18, only in C57BL/6J (C57BL/6J, ***p<0.001, P14 v/s P18) and stabilization at P22. The adult CD-1 and crossbreds had approximately 40% more dopaminergic neurons than C57BL/6J (Table 1, ^$$$$^p<0.0001, C57BL/6J v/s CD-1, F1X1 & F1X2). (b) The nigral volume increased during development from P2 to adulthood across all the mice strains, with a peak at P14. Throughout development, C57BL/6J had much smaller nigra than CD-1 and crossbreds (^$$$^p<0.001, C57BL/6J v/s CD-1, F1X1 & F1X2). A significant fall in nigral volume was noted in C57BL/6J between P14 to P18 (C57BL/6J, ***p<0.001, P14 v/s P18). The nigral volume was significantly higher in CD-1 and the crossbreds and stabilized at P14 whereas in C57BL/6J it reduced till P18. Note that the CD-1 and crossbreds possessed significantly larger SNpc at adulthood compared to that of C57BL/6J ($$p<0.01, v/s CD-1, FIX1 and F1X2). (c) The dopaminergic neuronal density in the CD-1, F1X1 and F1X2 strains, remained comparable throughout development. Interestingly, the density also increased gradually to reach peak at P14 in C57BL/6J (C57BL/6J, **p<0.01 P6 v/s P10, **p<0.01 P10 v/s P14). A stability in number of neurons per unit area was acheived by P22 but was significantly lesser than F1X2 (Adult, ^$$^p<0.01, C57BL/6J v/s F1X2).

**Table 1:**
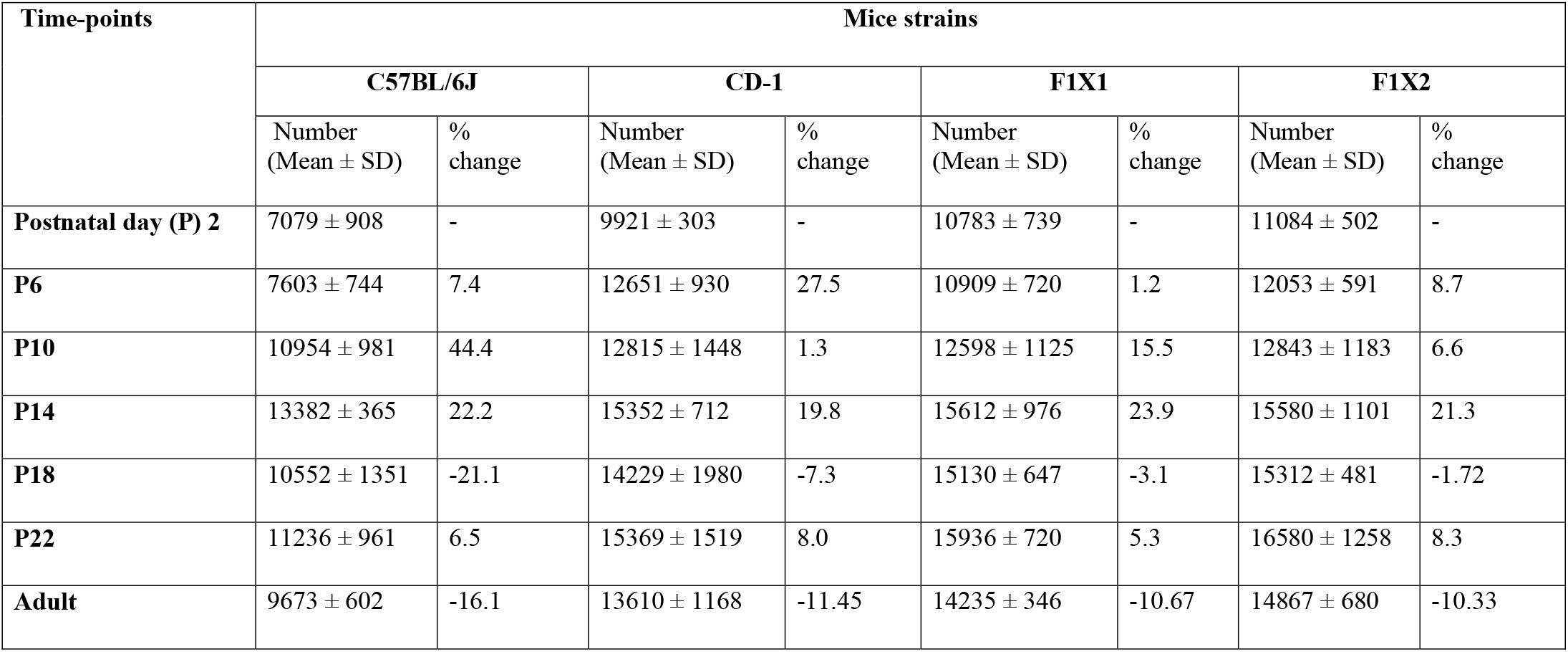
Table listing the number of dopaminergic neurons at each stage and the % difference. */**/***Significant alterations and the critical developmental time points. *p < 0.05

**Table 2:** Table listing the volume of substantia nigra pars compacta at each stage and the % difference in change. *Significant change denoting the critical developmental time points. *p < 0.05

| Time-points | Mice strains |  |  |  |  |  |  |  |
| --- | --- | --- | --- | --- | --- | --- | --- | --- |
|  | C57BL/6J |  | CD-1 |  | F1X1 |  | F1X2 |  |
|  | Volume (mm<sup>3</sup>)<br>(Mean ± SD) | %<br>change | Volume (mm<sup>3</sup>)<br>(Mean ± SD) | %<br>change | Volume (mm<sup>3</sup>)<br>(Mean ± SD) | %<br>change | Volume (mm<sup>3</sup>)<br>(Mean ± SD) | %<br>change |
| Postnatal day (P)2 | 0.22± 0.03 | - | 0.29± 0.003 | - | 0.30± 0.01 | - | 0.29±0.003 | - |
| P6 | 0.24± 0.02 | 11.79 | 0.37±0.02 | 27.35* | 0.31±0.01 | 5.49 | 0.32±0.01 | 9.11 |
| P10 | 0.29±0.02 | 20.95 | 0.33±0.03 | -9.13 | 0.36±0.03 | 15.89 | 0.35±0.05 | 11.00 |
| P14 | 0.30± 0.02 | 3.56 | 0.37±0.02 | 9.81 | 0.41±0.02 | 12.19 | 0.41±0.05 | 15.88 |
| P18 | 0.29± 0.03 | -4.86 | 0.37±0.03 | 1.84 | 0.40±0.02 | -2.38 | 0.37±0.02 | -8.12 |
| P22 | 0.34± 0.06 | 18.17 | 0.36± 0.08 | -2.35 | 0.41±0.02 | 3.98 | 0.41± 0.04 | 8.48 |
| Adulthood | 0.30± 0.02 | -11.53 | 0.39± 0.06 | 8.18 | 0.40± 0.06 | -4.05 | 0.37± 0.02 | -9.86 |

Thereafter a drastic fall in numbers was noted till P18, selectively in C57BL/6J (Fig. 2a, C57BL/6J, ***p<0.001, P14 v/s P18). The reduction stabilized at P22 (Fig. 2a, C57BL/6J), and the numbers were comparable with the adult mice described earlier (Vidyadhara et al., 2017). Despite the reduction between P14-P18, the numbers at P22 were significantly higher than those at P2. Interestingly, the CD-1 and crossbreds retained their numbers from P14 to stabilize with relatively more DA neurons at P18 and P22 (Fig. 2a). At P22, the CD-1 and crossbreds had approximately 40% more DA neurons than C57BL/6J (Fig. 2a, ^$$$$^p<0.0001, C57BL/6J v/s CD-1, F1X1 & F1X2). Effectively the quantitative differences between the strains persisted throughout the development. (Fig. 2a, Table 1, Adult, ^$$$$^p<0.0001, C57BL/6J v/s CD-1, F1X1 & F1X2).

### Alterations in nigral volume

Evaluation of nigral volume revealed two distinct observations. First that, a notable increase occurs in the nigral volume during development i.e. from P2 to P22 across all the mice strains (Fig. 2b). Secondly, at P2 and throughout development, the SN of C57BL/6J was much smaller than CD-1 and the crossbreds (Fig. 2b, ^$$$^p<0.001, C57BL/6J v/s CD-1, F1X1 & F1X2; differences at P2: C57BL/6J v/s CD-1: 40.15 %; v/s F1X1: 52.32 %; v/s F1X2: 56.58 %). A significant fall in nigral volume between P14 to P18 noted in C57BL/6J, paralleled a drastic reduction in the number of DA neurons (Fig. 2b, C57BL/6J, ***p<0.001, P14 v/s P18). Thus the nigral volume was significantly higher in CD-1 and crossbreds and stabilized at P14, whereas in C57BL/6J it reduced till P18 and stabilized thereafter (Fig. 2b). Essentially the CD-1 and crossbreds possessed significantly voluminous SN at P22 than the C57BL/6J (Fig. 2b, ^$$^p<0.01, v/s CD-1, FIX1 and F1X2).

### Differences in ‘density per unit area’ of DA neurons

The DA neuronal density in the CD-1, F1X1 and F1X2 strains, remained unchanged throughout development (Fig. 2c; Table 3). Interestingly, in the C57BL/6J, the density increased gradually to peak at P14 (Fig. 2c, C57BL/6J, **p<0.01 P6 v/s P10, **p<0.01 P10 v/s P14; Table 3) and stabilized by P22 but was significantly lesser than F1X2 (Fig. 2C, P22, ^$$^p<0.01, C57BL/6J v/s F1X2). Yet it was comparable to other strains.

**Table 3:** Table listing the tyrosine hydroxylase expression (staining intensity on a scale of 0-255) at each stage and the % difference in change.

| Time-points | Mice strains |  |  |  |  |  |  |  |
| --- | --- | --- | --- | --- | --- | --- | --- | --- |
|  | C57BL/6J |  | CD-1 |  | F1X1 |  | F1X2 |  |
| Postnatal day (P) 2 | Intensity (Mean ± SD) | % change | Intensity (Mean ± SD) | % change | Intensity (Mean ± SD) | % change | Intensity (Mean ± SD) | % change |
| P2 | 191.39 ± 6.74 | - | 218.22 ± 5.31 | - | 185.44 ± 8.43 | - | 211.11 ± 3.68 | - |
| P6 | 185.99 ± 8.14 | -2.82 | 204.65 ± 13.58 | -6.22 | 196.32 ± 2.34 | 5.86 | 195.28 ± 3.12 | -7.50 |
| P10 | 163.33 ± 11.17 | -12.18** | 193.24 ± 11.13 | -5.58 | 190.79 ± 7.63 | -2.82 | 186.98 ± 11.43 | -4.25 |
| P14 | 145.92 ± 4.97 | -10.66* | 152.70 ± 10.00 | -20.98***** | 171.88 ± 5.20 | -9.91* | 172.84 ± 6.30 | -7.56 |
| P18 | 158.73 ± 11.59 | 8.78 | 143.93 ± 5.86 | -5.75 | 157.23 ± 3.52 | -8.53 | 164.02 ± 2.28 | -5.10 |
| P22 | 133.23 ± 12.26 | -16.06*** | 159.64 ± 7.85 | 10.92 | 160.56 ± 1.74 | 2.12 | 162.77 ± 8.19 | -0.76 |
| Adulthood | 134.21 ± 2.63 | 0.73 | 146.61 ± 8.91 | -8.16 | 149.70 ± 3.08 | -6.76 | 161.52 ± 3.18 | -0.77 |

\*Significant change denoting the critical developmental time points. \*p < 0.05; \*\*\*\*\*p<0.0001, \*\*\*p<0.001, \*\*p<0.01, \*p<0.05

**Table 4:**
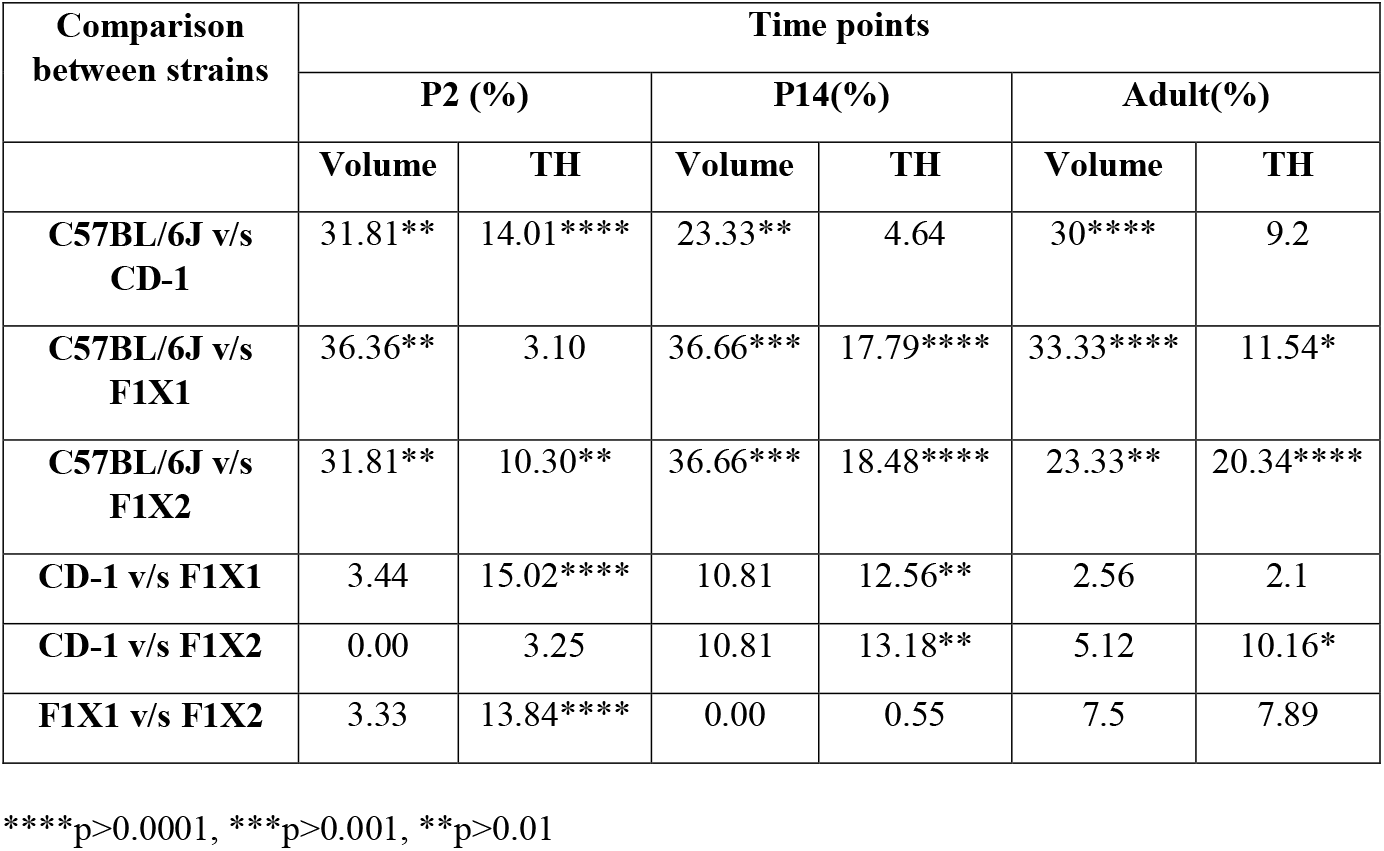
Comparison and percentage differences in volume of SN and tyrosine hydroxylase (TH) expression in different strains at critical time

### TH expression during development and Cellular morphology

Across all strains, the DA neurons increased in size with developmental age (Fig. 3a1-d5; e). A closer comparison revealed that, the P2 neurons of both the parent strains, i.e., C57BL/6J and CD-1 were much larger than the crossbreds (Fig. 3, compare a1, b1 with c1 & d1; P2, ^$$^p<0.01, C57BL/6J & CD-1 v/s F1X1 & F1X2). All the strains showed considerable increase in the soma size from P2-P10 (Fig. 3e, C57BL/6J, **p<0.01, P2 v/s P10; F1X1, ^###^p<0.001, F1X2, ^@@@@^p<0.001,P2 v/s P10). Interestingly, the soma sizes were comparable at P10, in all the strains, with a clear demarcation between the nucleus and cytoplasm. Thereafter, in C57BL/6J they continued to increase in size (Fig. 3e, C57BL/6J, **p<0.01, P18 v/s P22) resulting in a larger soma than the rest, at P22 (Fig. 3e, ^$$$^p<0.001, C57BL/6J v/s CD-1, F1X1 & F1X2).

**Fig. 3:**
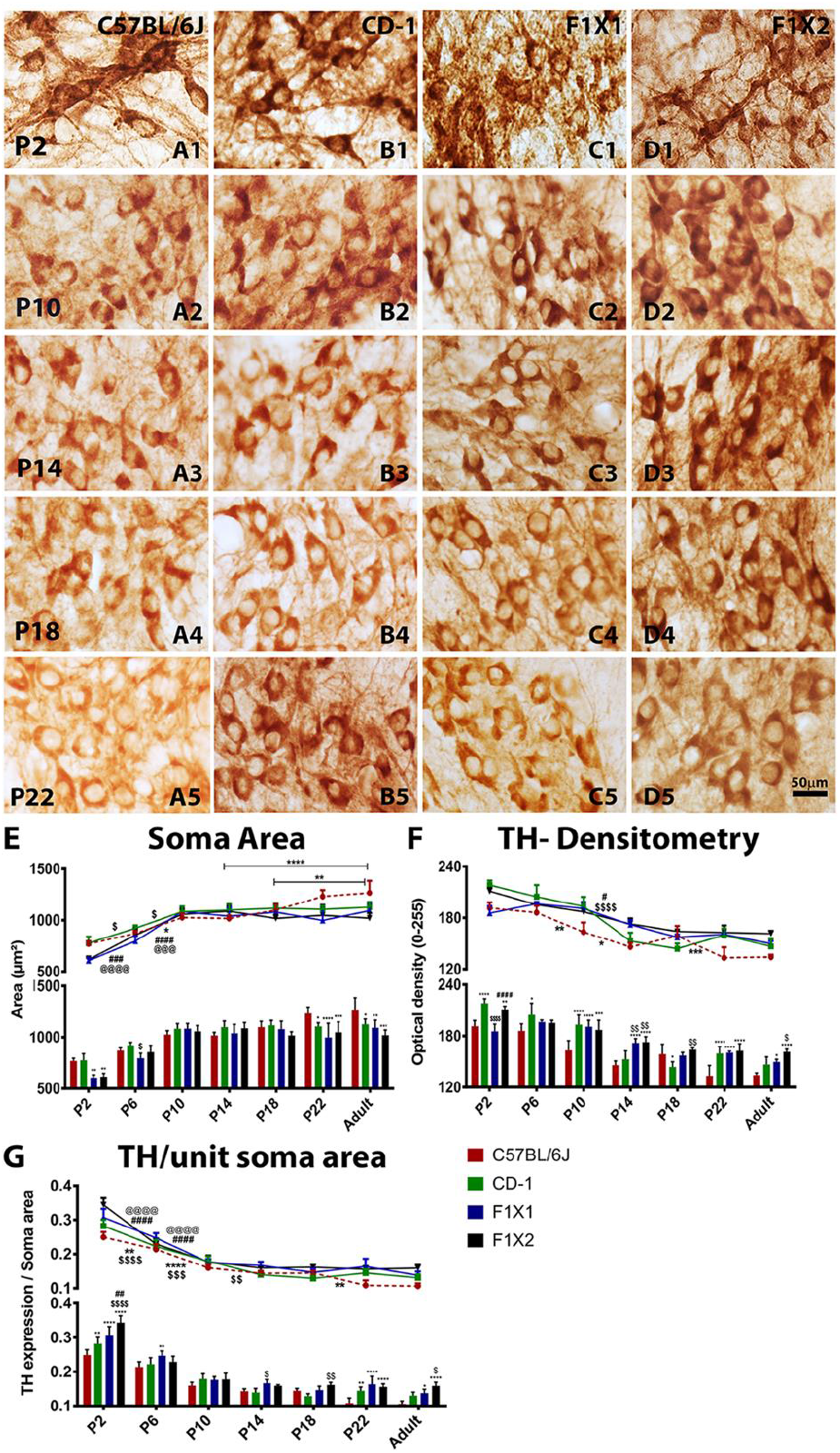
Representative photomigrographs of TH immunoractive SNpc neurons depicting the cellular morphology (a1-d5) and histograms (e-g) showing changes in size, cellular TH expression and TH/soma area during development: **Cellular size and morphology (e):** The dopaminergic neurons increased in size across strains (a1-d5) during development. At P2 the neurons of C57BL/6J and CD-1 were much larger than those of the crossbreds (a1 & b1 v/s c1 & d1 ^$$^p<0.01, C57BL/6J & CD-1 v/s F1X1 & F1X2). The soma size was comparable across strains at P10, when they attained adult phenotype i.e. clear nucleus and cytoplasm (a), the neurons of C57BL/6J mice continued to increase in size till P22 (a&e, C57BL/6J, **p<0.01, P18 v/s P22) resulting in significantly larger neurons at adulthood (a&e, ^$$$^p<0.001, C57BL/6J v/s CD-1, F1X1 & F1X2). **Cellular TH expression (f):** At P2 the TH expression in CD-1 and F1X2 was higher compared to C57BL/6J and F1X1. Stabilization of TH expression to its adult levels was noted from P14 to P18 in CD-1 and the crossbreds (Figure a1-d5; f), whereas a further reduction was seen in C57BL/6J which stabilized by P22 (f, C57BL/6J, ***p<0.001, P18 v/s P22). Although adult CD-1 and F1X1 showed a trend of higher TH expression, F1X2 expressed significantly higher TH than that of C57BL/6J (Adult, *p<0.05, C57BL/6J v/s F1X2). **TH expression per unit soma area (g):** The F1X2 mice showed significantly more TH expression per unit soma area at P2 than C57BL/6J (P2, ^$$$$^p<0.0001, C57BL/6J v/s F1X2). The reduction in the ratio stabilized at P14 in the CD-1, F1X1 and F1X2 midbrains, but at P22 in the C57BL/6J mice (C57BL/6J, **p<0.01, P18 v/s P22). The TH levels in nigral neurons of adult C57BL/6J were lower than other strains, especially when compared to F1X2 (Adult, ^$$^p<0.01, C57BL/6J v/s F1X2).

The CD-1 and F1X2 showed relatively higher TH expression at P2 (Fig. 3f). Interestingly, unlike other observations, it was highest at P2 compared to the other stages (Fig. 3f). The expression stabilized to its adult levels at P14 in CD-1 and crossbreds (Fig. 3f), while at P22 in C57BL/6J (Fig. 3f, C57BL/6J, ***p<0.001, P18 v/s P22).

In view of the appreciable differences in the soma size and the TH expression amongst strains, we quantified TH expression per unit soma area. The F1X2 mice nigra showed higher TH expression/unit soma area at P2 compared to C57BL/6J (Fig. 3g, P2, ^$$$$^p<0.0001, C57BL/6J v/s F1X2). A significant decline in the ratio was noted until P10 in all the strains. It stabilized at P14 in CD-1, F1X1 and F1X2, whereas at P22 in C57BL/6J with a further decrement (Fig. 3g, C57BL/6J, **p<0.01, P18 v/s P22).

### Alterations in developmental apoptosis and cell death markers

Developmental cell death was correspondingly evaluated, by TUNEL, at P2, P6, P10, P14, P18 and P22. TH immunoreactive (brown) cells labelled for TUNEL reaction (black) in the nucleus were identified as apoptotic DA neurons (Figure 4. Ab). TUNEL-ir cells that were TH negative, but present within the nigral region were also considered (Figure 4. Aa). A positive control (Figure 4. Ac) showed TUNEL reaction in almost all cells in the field. Peak apoptosis was noted at P2 and P14 (Figure 4B & 4C, P14 *p<0.05 C57BL/6J vs. F1X2) across the strains. The C57BL/6J had more TUNEL-TH co-labelled neurons, and persisted till P22; while in the CD-1 and the crossbreds, they subsided by P14.

**Figure 4:**
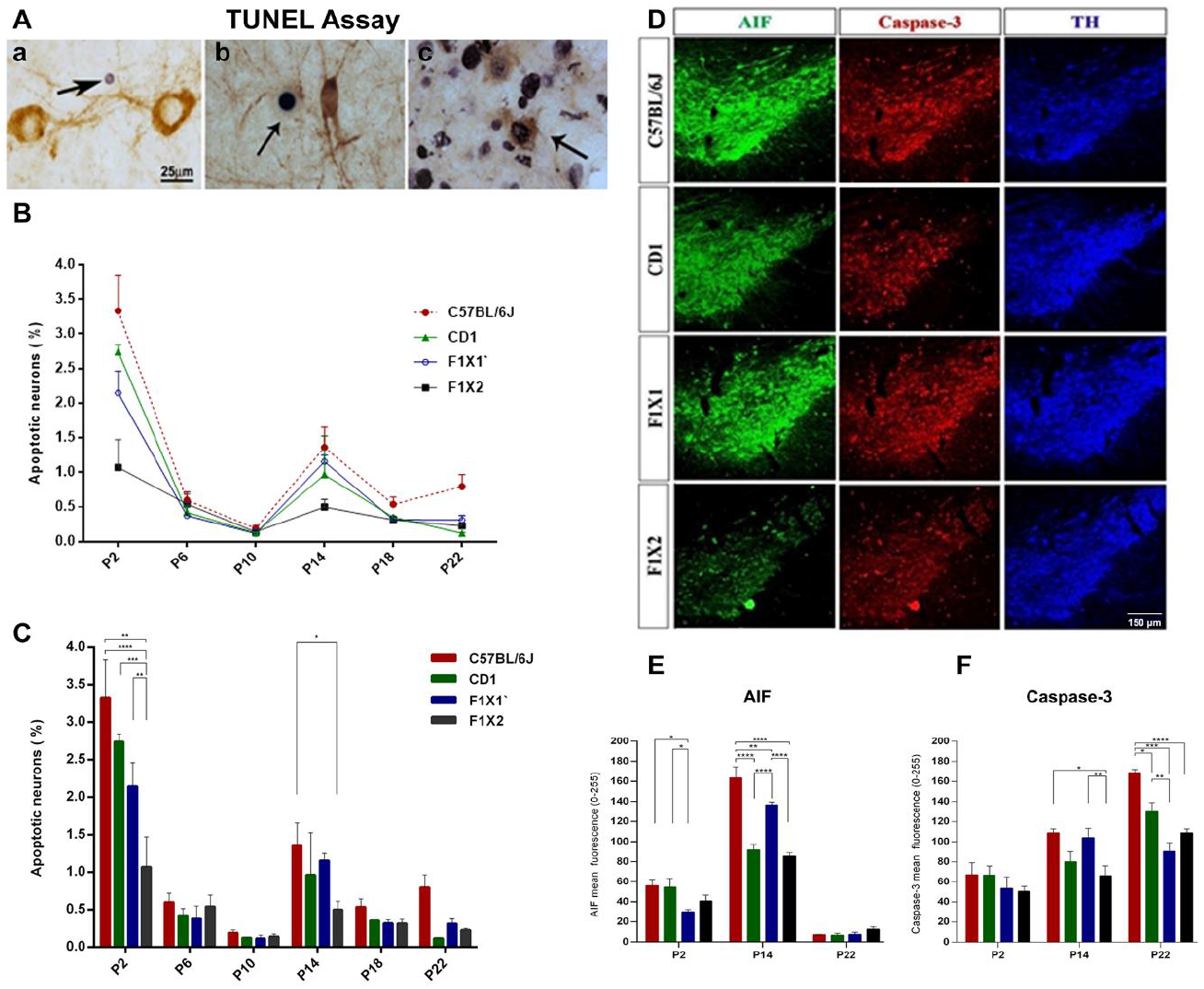
Relatively pronounced developmental apoptosis in MPTP-susceptible C57BL/6J mice nigra. A. Representative photomicrographs showing TUNEL-positive apoptotic cells (black) present within TH-ir DA neurons in the substantia nigra of C57BL/6J mice at P14. (Aa) TUNEL-ir and TH-ve and (Ab) TUNEL-ir and TH co-labelled were seen n nigra. (Ac). Positive control. B. Scale: 25 μm. Graph showing two peaks in apoptosis, one at P2 and another at P14 across all the strains during development, indicated by % apoptotic neurons. C. Histograms depicting %TUNEL-ir cells. Note the significantly higher % of TUNEL-ir cells in C57BL/6J at P2, than F1X1 and F1X2 (C, P2, C57BL/6J, **p<0.01 vs F1X1, ****p<0.0001 vs F1X2). CD-1 also had more apoptotic cells compared to F1X2 (C, P2, ***p<0.001, CD-1 vs F1X2). The difference in percent apoptotic neurons across mice strains was negligible at P6 and P10 (C, P6 and P10). At the second peak of P14, F1X2 had significantly fewer apoptotic neurons compared to C57BL/6J (C, P14, *p<0.05, C57BL/6J vs F1X2). D. Representative photomicrographs showing expression pattern of pro-apoptotic proteins AIF and caspase-3, co-labelled with TH in substantia nigra of C57BL/6J, CD-1, F1X1 and F1X2, at P14. (E & F) Histograms depicting immunofluorescence intensity measurements. E. Note, at P2, DA neuronal (TH +ve) AIF expression was higher in both the parent strains, especially when compared to F1X1 (E, P2, *p<0.05; C57BL/6J vs F1X1 and CD-1 vs F1X1). At P14, C57BL/6J showed a significantly higher AIF expression levels compared to CD1 and the crossbreds (E, P14, C57BL/6J, ****p<0.0001 vs CD-1, **p<0.01 vs F1X1, ****p<0.0001 vs F1X2). At P22, AIF expression was negligible across all the strains. F. Note, caspase-3 expression remained comparable across strains at P2. At P14, C57BL/6J showed a significantly higher caspase-3 expression compared to F1X2 (F, P14, *p<0.05, C57BL/6J vs F1X2, and a trend towards higher expression compared to CD-1. Note, caspase-3 expression in F1X1 was also significantly higher when compared to F1X2 (F, P14, **p<0.01, F1X1 vs F1X2). At P22, note a significantly higher caspase-3 expression in C57BL/6J when compared to CD1 and the crossbreds (F, P22, C57BL/6J, *p<0.05 vs CD-1, ***p<0.001 vs F1X1, ****p<0.0001 vs F1X2). Interestingly, caspase-3 expression was significantly lower even in CD-1 when compared to F1X1. CD-1 and F1X2 compared well (P22, **p<0.01, CD-1 vs F1X1). All photomicrographs were captured at 10X magnification.

The caspase-3 and AIF expression was evaluated exclusively in TH co-labelled neurons so as to assess the alterations in the surviving dopaminergic neurons. At P2, AIF expression was higher in both the parent strains (Figure 4E, *P<0.05; C57BL/6J vs F1X1 and CD-1 vs F1X1). At P14, it was much higher in C57BL/6J (Figure 4 D & E in comparison to F1X1 (**p< 0.01), CD-1 (****p<0.001) and F1X2 (****p<0.001).

Caspase-3 expression was comparable in all the study groups at P2. The differences began to be appreciable at P14 (Figure 4D) when it was higher in C57BL/6J (Figure 4F; *p <0.05 C57BL/6J vs. F1X2; *p <0.05 CD-1 vs. F1X2) as also in F1X1 (**p<0.01 vs F1X2) that persisted till P22 in C57BL/6J (C57BL/6J vs CD-1, *p<0.05; vs F1X1, ***p<0.001; vs F1X2 ****p<0.0001). In addition, at P22, the expression was also high in CD-1 (CD-1 vs F1X1, **p<0.01). Thus, maximum AIF expression was noted at P14 (Figure 4E) whereas peak caspase-3 expression was noted at P22 in C57BL/6J nigra (Figure 4F).

### GDNF expression in striatum and nigra

A significant upregulation in GDNF expression was noted in C57BL/6J between postnatal days 2 and 14, in the striatum (Figure 5. A & B; P2 *p<0.05 C57BL/6J vs. CD-1; P2 **p< 0.01 C57BL/6J vs. F1X2; C57BL/6J **p< 0.01 P2 vs. P14; P14 *p< 0.05 C57BL/6J vs. F1X1). The basal GDNF was high in CD-1 and F1X2 at P2, and also at P14. In contrast, in the nigra, its expression reduced between P2 and P14 in all the strains. (Figure. 5C & D. P2 **p< 0.01 CD-1 vs F1X2; P2 *p< 0.05 F1X1 vs. F1X2; C57BL/6J ***p< 0.001 P2 vs. P14; CD-1**p< 0.01 P2 vs. P14; F1X2 *p< 0.05 P2 vs P14).

**Figure 5:**
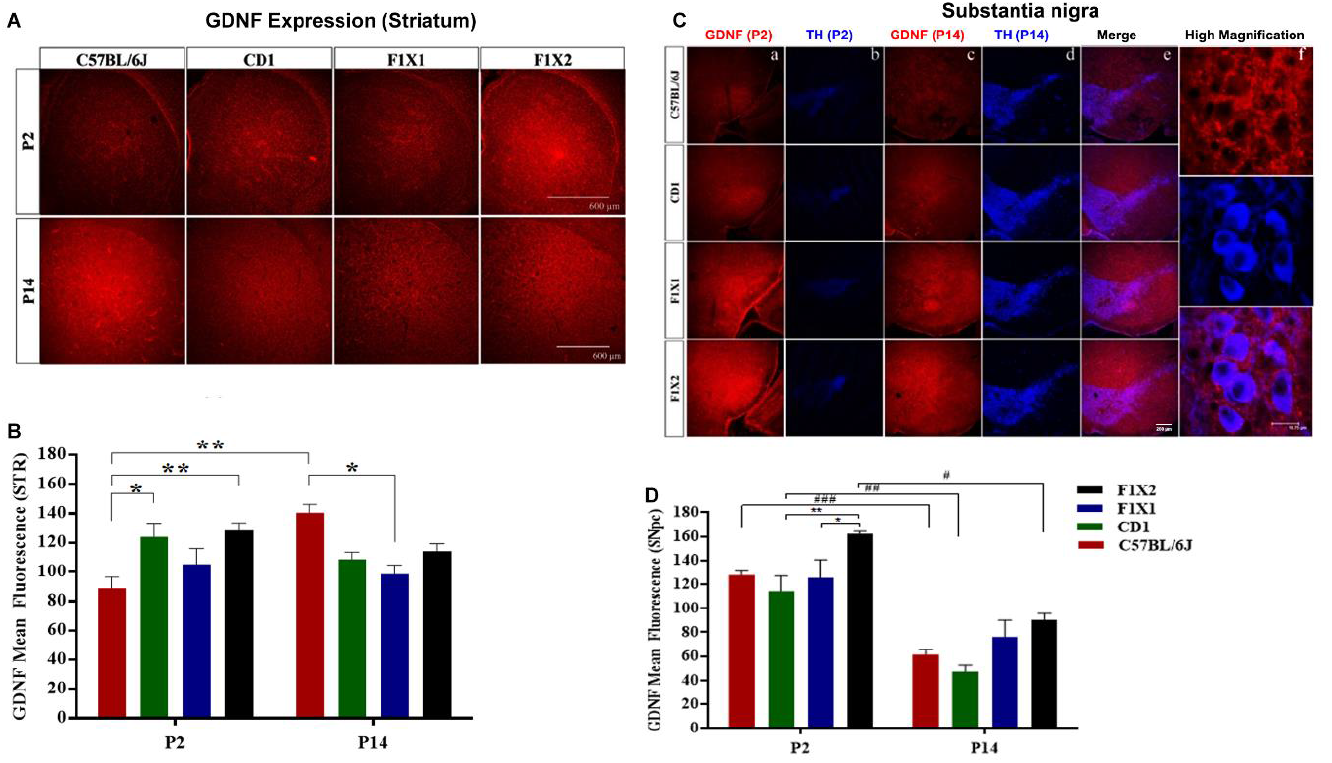
Nigrostriatal GDNF expression at time-points of peak apoptosis. A. Representative photomicrographs of striatal GDNF expression in C57BL/6J, CD-1, F1X1 and F1X2, at P2 and P14 time-points. B. Histograms depicting immunofluorescence intensity measurements. Note, striatum of C57BL/6J has significantly lower GDNF levels compared to CD-1 and F1X2 at P2 (B, P2, C57BL/6J, *p<0.05 vs CD-1, **p<0.01 vs F1X2). In C57BL/6J compared to P2, it was more at P14 (B, C57BL/6J, **p<0.01, P2 vs P14). At P14 the GDNF expression was significantly higher in C57BL/6J than F1X1 (B, P14, *p<0.05, C57BL/6J vs F1X1). Scale: 600 μm. C. Representative photomicrographs of GDNF and TH co-labelled substantia nigra of C57BL/6J, CD-1, F1X1 and F1X2, at P2 and P14. Histogram (D). Note a significantly higher level of GDNF at P2, in F1X2 (D, P2, F1X2, **p<0.01 vs CD-1, *p<0.05 vs F1X1). At P14, both F1X1 and F1X2 showed moderately higher nigral GDNF expression compared to parent strains (D, not significant). Note a significant down-regulation of nigral GDNF from P2 to P14 in all the strains, except F1X1 (D, P14, C57BL/6J, ^###^p<0.001 P2 vs P14, CD-1, ^##^p<0.01 P2 vs P14, F1X2, ^#^p<0.05 P2 vs P14). Scale: enlarged: 18.75 μm. All low magnification images were captured using 10X objective.

## Discussion

A few neuropsychiatric diseases like autism and attention deficit hyperactive disorder etc. are thought to be neurodevelopmental disorders while others like Huntington’s, Alzheimer’s, Parkinson’s disease are designated as adult or elderly onset disorders (Lee *et al.* 2012; Panzer & Viljoen 2005; Fatemi & Folsom 2009; Sullivan & Brake 2003; Weber *et al.* 2008). It is being increasingly recognized that deregulation of developmental neuronal apoptosis or glial death may result in functional deficits and associated diseases in the later years of life (Arya and White, 2015; Lein et al., 2018). In the absence of human tissues for analysis, animal models are valuable resources for mechanistic explanations of a disease. Ours is the first study using unbiased stereology on developing nigra that connects the developmental trajectories of nigral DA neurons and susceptibility of mice to MPTP. The studies were conducted in MPTP-sensitive C57BL/6J, MPTP resistant CD-1 mice and the F1 generation of their reciprocal crossbreds. Our model offers the advantage of “no use of neurotoxins”, since most other studies on PCD relied on chemical lesions to investigate target dependence in development.

The first two postnatal weeks are crucial for the rodent midbrain DA system, for being associated with culmination of neuronal migration, maturation of perikarya, extension of axons and their terminal differentiation; which occurs alongside the apoptotic death of neurons that fail to mature (Voorn *et al.* 1988; Oo & Burke 1997; Antonopoulos *et al.* 2002; Burke 2003). Stressors during this period prime the DA system to subsequent insults like prenatal stress, maternal separation, ischemia, hypoxia and excitotoxicity etc. resulting in the delayed attainment of DA phenotype (Katunar *et al.* 2010), reduction in the number of DA neurons, exaggerated response to neurotoxins (Pienaar *et al.* 2008; Mabandla & Russell 2010) and affect the DA neurotransmission at adulthood (Rodrigues *et al.* 2011). In fetal non-human primates (Morrowa et al., 2007) peak PCD of DA neurons occurs at E80 and approximately at mid-gestation in humans, when approximately 50% neurons are pruned. The time period P2 in rats equates with E79/80 in monkeys (Clancy et al., 2001), thus P2 in mice is a critical window of vulnerability and protective modalities could be developed if the relevant molecular underpinnings are understood.

The clusters of TH-ir neurons found along the midline during early postnatal development i.e. P2 and P6 probably represent the migrating neurons. The gradual increase in numbers till P14 point at the culmination of neuronal migration (Park *et al.* 2000; Lieb *et al.* 1996; Kawano *et al.* 1995), or addition of TH-ir neurons by postnatal neurogenesis (Lie *et al.* 2002; Shan *et al.* 2006; Zhao *et al.* 2003; Park *et al.* 2000). The third likelihood is that progressively more neurons show detectable levels of TH expression i.e. they attain DA phenotype (Chocyk *et al.* 2011; Bjorklund & Dunnett 2007; Katunar *et al.* 2010). The delay in attainment of adult nigral architecture in C57BL/6J i.e. at P18, may be attributed to the ongoing cell loss vis-à-vis the early maturation at P14 in CD-1 and the crossbreds. Our observations on C57BL/6J, match those of Oo and Burke (1997), who showed two peaks of loss i.e. at P2 and P14, proposing a biphasic nature of programmed cell death. We propose an additional minor peak at P22 in C57BL/6J, complemented by TUNEL reaction, caspase-3 expression and reduction in TH-ir dopaminergic neurons.

Both caspase-mediated and caspase-independent cell death pathways, associated with mitochondria and endoplasmic reticulum respectively, work in unison to cause neurodegeneration (Sanges et al., 2006). Increased AIF expression during development corroborates with the role of caspase independent cell death in developing human midbrain (Pagida et al., 2020). High AIF expression at P14 in addition to peak caspase-3 expression at P22, corroborated by TUNEL-expression, further illustrate the alliance between the two cell death pathways in the instance of DA neurons.

Classical studies suggest that GDNF protects nigral neuron from apoptosis (Burke, 1998; Oo et al., 2003). Recent studies consider it as a potent therapeutic for PD (Cheng et al., 2018; Whone et al., 2019). It is expressed in the striatum and reaches SNpc through retrograde transport (Tomac et al., 1995). Therefore, its mRNA synthesis and protein expression during striatal development are critical (Blum & Weickert, 1995). The upregulation of GDNF in the striatum from day 2 to day 14, in contrast to reduced expression in the SNpc at analogous stages, compares well with earlier studies (Pedro et al., 2005; Blum & Weickert, 1995) and it may be associated with the refinement of the nigrostriatal connections. GDNF expression at P2 may be paracrine/nigral in origin, since the nigro-striatal connections are yet not formed. At later stages, the source may switch to striatum. The higher levels in the crossbreds suggest better neuroprotection in them, which could be conjectured during nigral development in the Anglo-Indians, scaling up their dopaminergic reserve. It might be worthwhile to examine the early embryonic periods and develop preventative strategies.

Cellular hypertrophy following second peak of apoptosis noted exclusively in C57BL/6J may be a physiological process corresponding to the expansion of nerve terminals, also seen in other amine systems (Park *et al.* 2000; Saito *et al.* 1996; Fujimiya *et al.* 1986b; Fujimiya *et al.* 1986a). Alternatively it may be a predisposing factor; since larger neurons are vulnerable to degeneration (Wang & Michaelis 2010). Hypertrophy of DA neurons in aging human nigra was considered as a compensation for sub-threshold neurodegeneration (Cabello et al, 2002) and its absence as a marker for resilience (Alladi et al., 2009). The gradual decrease in cellular TH over development alongside an increase in soma area, suggests that overall dopamine synthesis remains stable through development. Amongst the strains, F1X2 showed best preservation of TH levels.

A correlation between the nigral volume and the neuronal numbers through development and across strains indicates that volume can be an indirect measure of DA neuronal number. Nigral volumetry by MRI is a diagnostic tool for PD, based on a similar premise. The striatal volume varies in healthy individuals (Harris *et al.* 1999; Rosas *et al.* 2001) and imparts differential susceptibility to psychiatric disorders (Voelbel *et al.* 2006; Kreczmanski *et al.* 2007; Reiss *et al.* 1993). Anthropometric studies suggest that children with smaller head circumference and cerebral volume may develop Huntington’s disease (Lee *et al.* 2012) reflecting the developmental origin of the disease. Cognitive capabilities were better protected in Alzheimer’s disease patients with relatively larger head circumference (Perneczky *et al.* 2010). Therefore, nigral volumetry may be a valuable tool to identify the predisposed individuals/populations.

A hypothetical division of the postnatal period in three phases reveals that the second phase that defines the neuronal numbers is more critical. Such differences may be envisioned in the Caucasian, Asian-Indians, and their admixed population imparting them varying degrees of susceptibility to develop PD (Chu *et al.* 2002; Alladi *et al.* 2009; Das *et al.* 2010; Strickland & Bertoni 2004; Ragothaman *et al.* 2003; Alladi *et al.* 2010a; Alladi *et al.* 2010b).

Finally, the findings of fewer neurons throughout development in C57BL/6J nigra; posit the presence of excessive endogenous toxins or pro-apoptotic factors at these stages. This possibility was strengthened by corresponding high levels of caspase-3 and AIF. The higher cellular packing density at P14 in C57BL/6J may represent increased competition for trophic factors. Whereas, minimal loss of DA cells, higher GNDF and controlled expression of pro-apoptotic factors in CD-1 and the crossbreds, hint at better endogenous milieu. Developmental apoptosis sculpts aberrant neural connections (Kuan et al., 2000; Buss et al., 2006) to regulate the final neural numbers in adult multicellular organisms (Yamaguchi & Miura, 2015; Castagna et al., 2016). Since both the mice strains are non-transgenic and genetically distinct, we hypothesize that genetic constitution may predispose individuals or populations even in absence of major mutations. In late eighties, Fahn (1989) hypothesised that the brain wiring during development may influence the number of neurons at birth and other cellular features which may be unique to the individuals, making them either vulnerable or resilient to PD. He also proposed that the metabolic processes in the brain may govern vulnerability to develop PD (Fahn 1989). Both these hypotheses hereby stand validated, hinting at the developmental origin of vulnerability to MPTP. It may therefore be possible that susceptibility to PD originates during nigral development in humans.

## Acknowledgements

We are grateful to Dr. Bindu M. Kutty, Prof. and Head, Dept. of Neurophysiology, NIMHANS for laboratory facilities, Dr. G.H. Mohan, Head Veterinarian at NCBS, Bengaluru for providing breeding colonies of CD-1 mice strain. VDJ received Movement Disorder Society (MDS) International Congress travel grant to present part of the study at MDS Congress 2017, Vancouver, Canada. HY received Dept. of Biotechnology, Govt. of India travel award to present a part at MDS Congress, 2019.at Nice, France.

## Ethics approval

All applicable international, national, and/or institutional guidelines for the care and use of animals were followed. All the procedures performed in the studies involving animals were in accordance with the ethical standards of NIMHANS which adhere to the CPCSEA and NIH guidelines.

## Abbreviations

ABC: Avidin Biotin Complex
ANOVA: Analysis of Variance
AIF: Apoptosis Inducing Factor
CPCSEA: Committee for the Purpose of Control and Supervision of Experiments on Animals
GDNF: Glial Derived Neurotropic Factor
MPTP: 1-methyl-4-phenyl-1,2,3,6-tetrahydropyridine
Nurr1: Nuclear receptor related-1 protein
P: Postnatal day
PBST: Phosphate Buffered Saline Containing 0.1% Triton X-100
PD: Parkinson disease
PitX3: Paired Like Homeodomain 3
PCD: Programmed Cell Death
SEM: SD: Standard Deviation
SN: Substantia Nigra
SNpc: Substantia Nigra pars compacta
SNpr: Substantia Nigra pars reticulata
TUNEL: Terminal deoxynucleotidyl transferase (TdT) dUTP Nick-End Labeling
TH: Tyrosine Hydroxylase
TH-ir: TH-immunoreactive

